# Adipogenic differentiation of hematopoietic lineage cells isolated from adipose tissue of humans

**DOI:** 10.1101/2024.06.22.600230

**Authors:** Kathleen M. Gavin, Bogdan Conrad, Timothy Sullivan, Ruby Vianzon, Alistaire S. Acosta, Wendy M. Kohrt, Dwight J. Klemm

## Abstract

We previously discovered some adipocytes in the major white fat depots of mice and humans arise from bone marrow-derived cells of hematopoietic lineage rather than conventional mesenchymal precursors, termed bone marrow-derived adipocytes (BMDA). Here we aimed to determine if hematopoietic lineage cells isolated from adipose tissue and circulation of humans could undergo adipogenic differentiation *in vitro,* thereby establishing an *in vitro* model for studies of BMDA. We hypothesized that hematopoietic lineage cells isolated from adipose tissue, but not circulation, of humans would demonstrate adipogenic potential. Participants included younger (20-50 years) and older (>50-75 years) men and women, BMI 20-37 kg/m^2^. Subcutaneous abdominal adipose tissue biopsies were obtained and stromal cell populations identified by flow cytometry. Sorted cells underwent *in vitro* cultivation via traditional mesenchymal culture methodology (mesenchymal lineage) or a novel 3D-fibrin clot followed by traditional adherent culture (hematopoietic lineage) for assessment of proliferation and differentiation capacity. We found hematopoietic lineage cells isolated from the adipose tissue stroma, but not the circulation, were capable of proliferation and multilineage (adipogenic and osteogenic) differentiation *in vitro*. We provide a new investigative tool that can be used to perform translational studies of BMDAs and provide initial evidence that hematopoietic lineage cells isolated from the adipose tissue of humans can undergo hematopoietic-to-mesenchymal transition with multilineage differentiation potential in an *in vitro* environment.

## Introduction

Until recently, the complexity of the cellular composition of adipose tissue has been underappreciated. Although by volume adipocytes are the primary component of adipose tissue, there are a myriad of other cell types that constitute the stromal-vascular (non-adipocyte) fraction (SVF).(1) Undoubtedly, the characteristics of adipocytes themselves are a critically important factor in determining the overall metabolic phenotype of a given adipose tissue depot.(1) We now recognize that more than one ‘type’ of adipocyte can populate a single fat depot (2, 3) and evidence continues to mount that adipocyte origins differ between and even within adipose tissue depots.(4–6)

Adipocytes turn over slowly, with only about 10% of cells replaced each year in humans, even during weight stability.(7) Thus, it is essential that a pool of progenitor cells are present within the adipose tissue, ready to undergo commitment to the adipocyte lineage. For years, dogma held that all new adipocytes arise from tissue-resident stem cells of the mesenchymal lineage.(8) However, immune cells of the hematopoietic lineage are also located in adipose tissue.(9) The contribution of these cells to adipogenesis was long dismissed until recent studies in mice and humans demonstrated that cells of the hematopoietic lineage contribute to adipogenesis in major white fat depots.(6, 10–15) These bone marrow-derived adipocytes (BMDAs) can make up from <5-35% of cells in a given adipose tissue depot, levels that could influence physiology.(6, 11, 14)

Murine studies demonstrate that upon mesenchymal transition, cells of the hematopoietic lineage lose their hematopoietic-specific surface markers.(13) However, an alternative unique biomarker to distinguish BMDAs from conventional adipocytes has not yet been discovered. Thus, identification and study of mature BMDAs has been limited to murine lineage tracing strategies and DNA chimerism analysis in patients having had a bone marrow or hematopoietic stem cell transplantation for clinical indication. Here we describe a human primary BMDA cell culture model that can be utilized to perform *in vitro* studies of BMDAs until additional clinical identification tools are available.

The aim of this study was to determine if hematopoietic lineage cells isolated from the adipose tissue and circulation of humans could undergo adipogenic differentiation *in vitro*. A secondary outcome was to determine if there were sex or age differences in the presence of conventional mesenchymal or hematopoietic lineage adipocyte precursors in subcutaneous abdominal adipose tissue. We hypothesized that hematopoietic lineage cells isolated from the adipose tissue, but not the circulation, of humans would demonstrate adipogenic potential.

## Materials and Methods

### Study Participants

The study was approved by the Colorado Multiple Institutional Review Board and written informed consent was obtained from all participants. All procedures took place at the Clinical and Translational Research Center (CTRC) on the University of Colorado Anschutz Medical Campus (CU-AMC). Samples were obtained from participants enrolled in NCT02654925, NCT02758431, and NCT04043520.

Participants were healthy males and females either 20-50 or >50-75 years of age (y). Older women were postmenopausal as identified by no menstrual cycles within the last 12 months and confirmed by follicle-stimulating hormone (FSH) ≥ 30 mIU/mL. Premenopausal women were not currently pregnant or lactating. Inclusion criteria included a BMI between 20 and 40 kg/m^2^. Exclusion criteria included current (within the last 6 months) use of any hormone replacement or hormonal contraceptive, use of glucose lowering medication and diagnosis of type 2 diabetes or uncontrolled metabolic disorders.

### Body composition

Body composition was measured by dual x-ray absorptiometry (DXA) using a Hologic Discovery-W instrument (software Apex 4.0.1; Hologic, Waltham, MA). Total body scans were analyzed for total and regional fat mass, and fat-free mass (FFM).

### Blood draw and circulating factors

Blood samples were obtained through an antecubital blood draw after ≥8 hour fast. After collection, samples to be analyzed for circulating factors were stored at -80°C until batched analysis. Blood samples for isolation of circulating hematopoietic cells were collected into EDTA tubes and kept on ice until processed simultaneously with adipose tissue samples.

All assays were Beckman Coulter, performed by the CTRC Core Laboratory. Estradiol (E2) and FSH were measured by chemiluminescent immunoassay, Testosterone via 1-step competitive assay, glucose and insulin by hexokinase, UV and chemiluminescent immunoassay.

### Adipose tissue biopsy collection and processing

The skin and subcutaneous abdominal adipose tissue directly adjacent to the umbilicus underwent local numbing with 1% lidocaine without epinephrine, after which the adipose tissue samples were obtained using a modified Coleman’s manual vacuum (“mini liposuction”) technique. A small incision was made, followed by infiltration of the adipose tissue with ∼30-50mL of 0.15% tumescent lidocaine solution in 0.9% normal saline using a Coleman infiltration cannula. Approximately one to three grams of adipose tissue was removed using a Coleman aspiration cannula.

Adipose tissue was rinsed with collection buffer (Krebs-Ringers-HEPES (KRH), 2.5mM glucose, 200nM adenosine, pH 7.4) and obvious blood vessels and clots removed. 250-500mg was flash frozen in liquid nitrogen and stored at -80°C. Remaining tissue was transported to the laboratory where collection buffer was replaced with prewarmed digestion buffer (KRH, 2.5mM glucose, 2% fetal bovine serum [FBS], 200nM adenosine, 1mg collagenase/0.25g tissue, pH 7.4). Tissue was digested for 30-60 minutes in a shaking water bath maintained at 37°C. The resultant cell suspension was filtered through 250μm mesh and digestion stopped with 1X volume of wash buffer (Hanks Balanced Salt Solution [HBSS] + 2% FBS + 200nM adenosine). Samples were spun at 300g x 10 min, the floating adipocytes removed to a clean tube, and wash steps repeated. The stromal pellets were combined and spun at 500g for 10 min to pellet the cells. Here forward, all spins were at 500g x 10 min unless otherwise stated. The stromal cell pellet was resuspended in 10mL Erythrocyte Lysis Buffer and incubated for 10 min on ice. Simultaneously, 1mL of blood underwent red blood cell lysis. When applicable, adipose and blood samples were processed in parallel from here forward. After erythrocyte lysis, samples were spun and cell pellet resuspended in 1mL wash buffer. Live cells were counted via trypan blue detection on a Cellometer X1 (Nexcelom Bioscience LLC, Lawrence, MA) cell counter.

### Flow cytometry

After counting, cells were spun and resuspended at a concentration of 1x10^6^ cells/100μL in wash buffer. Human TruStain FcX was added at 5μL/100μL cell suspension and cells incubated on ice for 10 min. All lineage antibodies were added at 5μL per million cells in 100µL staining volume (or 0.25μg/10^6^ cells) and mixed gently. Samples were incubated on ice in the dark for 25 min. Following incubation, samples were spun, the supernatant aspirated, and the cell pellet resuspended in wash buffer. After removal of the second supernatant, the cells were resuspended in flow buffer (HBSS + 5% FBS + 200nM adenosine, pH 7.4) with volumes adjusted to 500μL – 1mL for cell sorting. DAPI was added to a final concentration of 1mg/mL. Cell sorting was initiated within 15 minutes. Unstained cells, single color compensation beads and fluorescent minus one (FMO) controls were used. Antibodies are listed in the supplementary information.

SVF cells were sorted using an Astrios cell sorter with Summit 6.3 software (Beckman Coulter, Fullerton, CA, USA). A 70µm nozzle tip was used with a sheath pressure of 60Ψ and a drop drive frequency of approximately 95,000 Hz with an amplitude of 15V. Manufacturer’s protocols were followed to quality control and set up the sorter. The sheath fluid was IsoFlow. The sample and collection tubes were maintained at 10°C using an attached recirculating water bath. To keep cells in suspension, the Astrios’ SmartSampler sample station was set to maintain an agitation cycle of 5 seconds on and 8 seconds off. The sample flow rate was set to a pressure differential of <1.0Ψ. Sort mode was set to Purify 1. Appropriate signal compensation was set using single-color control samples. The flow cytometry gating strategy is included in **Figure 1**. Cells were identified with the following surface markers: mesenchymal lineage cells, CD45^NEG^; conventional mesenchymal adipocyte precursor cells, CD45^NEG^/CD31^NEG^/CD29^POS^ (16); hematopoietic Lineage, CD45^POS^; myeloid lineage, CD14^POS^; macrophages (ATM) & monocytes, CD14^POS^/HLA-DR^POS^ with divergent expression of CD206 (ATM CD206^POS^, monocytes CD206^NEG^); neutrophils, CD14^POS^, CD16^POS^, HLA-DR^NEG^, Siglec-8^NEG^; basophils, CD14^NEG^/Siglec-8^POS^; lymphocytes, CD14^NEG^/CD206^NEG^. Additional analysis of the lymphocyte population identified both CD3^POS^/CD8^POS^ and CD3^POS^/CD4^POS^ T cells (data not shown).

**Figure 1.**
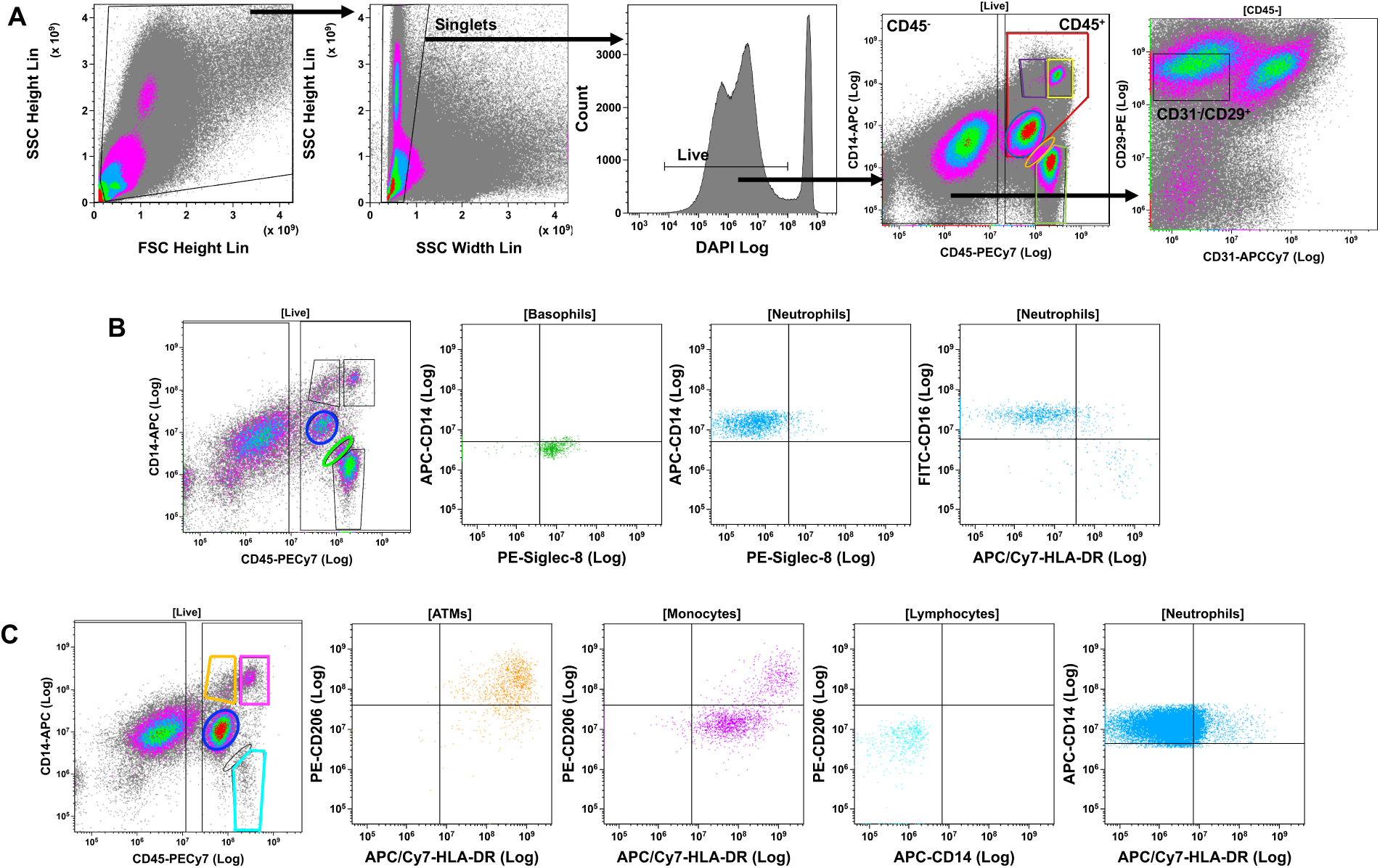
Flow Cytometry/Fluorescent Activated Cell Sorting (FACS) strategies for identifying mesenchymal and hematopoietic lineage cells isolated from stromal vascular fraction (SVF) of human adipose tissue samples. **(A)** SVF cells were stained with fluorescent antibodies and mesenchymal and hematopoietic subpopulations were purified by FACS using the representative gating strategy shown. Debris was excluded based on the ratio of forward scatter (FSC) height to side scatter (SSC) height. Clusters and aggregates were excluded based on the ratio of SSC width to SSC height. Live cells were DAPI negative. CD45 fluorescence revealed distinct positive and negative populations. The CD45^NEG^ population represented mesenchymal lineage cells, which included CD31^NEG^/CD29^POS^ conventional mesenchymal adipocyte precursor cells. The CD45^POS^ population contained multiple populations of hematopoietic lineage cells: red gate - myeloid lineage (CD14^POS^), including purple gate - macrophages (ATM), yellow gate – monocytes and blue gate – neutrophils; orange gate – basophils; green gate – lymphocytes (CD14^NEG^). **(B/C)** Representative images for identification of hematopoietic lineage subpopulations by flow cytometry from two different cell donors. The flow cytometry strategy was identical to the first four panels in A, with addition of fluorescent antibodies for the identification of hematopoietic subpopulations. **(B)** Basophils: CD14^NEG^/Siglec-8^POS^; Neutrophils: CD14^POS^, CD16^POS^, HLA-DR^NEG^, Siglec-8^NEG^. **(C)** ATM and monocytes: CD14^POS^/HLA-DR^POS^ with divergent expression of CD206 (ATM CD206^POS^, monocytes CD206^NEG^). Lymphocytes: CD14^NEG^/CD206^NEG^. Additional analysis of the lymphocyte population identified both CD3^POS^/CD8^POS^ and CD3^POS^/CD4^POS^ T cells (data not shown). ATM and monocyte populations were pooled for analysis because of difficulty with complete discrimination and low numbers for downstream analysis. (Use color in print version)

Circulating blood samples were obtained from a subset of participants (n=5 young males, n=3 young females, n=5 older males, n=4 older females) and analyzed by an identical flow cytometry strategy to adipose tissue SVF (**Supplementary Figure 1**).

All cells were sorted into growth medium (MEM α + 10% fetal bovine serum + 0.1x penicillin/streptomycin) and transported back to the lab on ice. Cell sorting was completed in the University of Colorado Cancer Center Flow Cytometry Shared Resource.

Purity of sorting was indicated at 97% for Neutrophils and 94% for Lymphocytes (**Supplementary Figure 2**). Because of the extremely low cell numbers for the ATM & monocyte populations, purity of that population was not independently evaluated.

### Cell culture

Sorted cells were gently vortexed and spun for 10 min at 500g. Cell pellets were resuspended in growth medium and conventional mesenchymal lineage cells were plated directly onto cell culture treated plastic plates for proliferation by standard techniques (17) at a density of ∼4,500 cells/cm^2^. Fresh growth media was applied the next day and media changes took place every 2-3 days throughout the studies. Cells were grown to 70-80% confluence, detached with 30-60 min room temperature incubation in Accutase and passaged 1-2 additional times. Some cells were then seeded immediately for experiments and remaining cells were frozen down in liquid nitrogen for use in future experiments.

Hematopoietic lineage cells were cultured according to previously published methods (13) with some modifications. ATM/monocytes, neutrophils, and lymphocytes were each resuspended in 10μL growth media at 40,000 cells/μL. Fibrinogen (60 μL, 5 mg/mL in 0.9% saline) was added, mixed gently and transferred to one well of a 96 well plate (per 400,000 cells) containing 1.4μL of thrombin (50 U/mL) and the suspension mixed by gentle pipetting. Clots formed over 30–60 minutes and growth media was subsequently overlaid. Media was refreshed every 2-3 days throughout the study.

Cells were recovered from the clots after 5-7 days. The culture medium was removed and matrices rimmed with a 27 gauge needle. The remaining liquid in the well was removed and 100μL of digestion mix (75μL growth medium + 23μL bovine plasminogen (1.12mg/mL) + 2μL urokinase (20U/μL)) was added for 60 minutes at 37°C, mixing by pipetting every 20 minutes. After digestion, cells were pelleted in a 1.5mL tube at 500g x 10 minutes, resuspended in growth medium, and replated directly onto plastic into the same number of wells of a 96 well plate. All cells then underwent identical culture methodology to the mesenchymal lineage cells.

### Cell viability (proliferation)

Cells used in all experiments were passage 3-5. 2,000 cells were seeded per well in 96 well plates; 3 replicates per subject/cell type were measured at 24, 48, and 72h. Absorbance, indicative of number of viable cells, was measured at 490nm after 4h incubation with CellTiter 96 AQueous One Solution according to the manufacturer’s protocol.

### Multilineage differentiation

*Adipogenesis.* Differentiation was induced according to the protocol published by Lee et. al.(17) In brief, after reaching 100% confluence, growth media was exchanged for complete adipocyte differentiation medium (CDM; Dulbecco’s Modification of Eagle’s Medium/Ham’s F-12 50/50 Mix [DMEM/F12], 33µM Biotin, 17µM pantothenate, 100nM dexamethasone, 1µM rosiglitazone, 0.5mM IBMX, 2nM T3, 10µg/mL transferrin, and 100nM insulin) for 7 days. This was followed by 7 days of adipocyte maintenance medium (AMM; DMEM/F12, 33µM Biotin, 17µM pantothenate, 10nM dexamethasone and 10nM insulin). Adipogenic potential was quantified by Oil Red O staining (see Supplementary Methods).

*Osteogenesis*. Osteogenic differentiation potential was assessed by seeding six wells of each cell type in a 24-well plate at 10,000 cells/well. Upon 80% confluency, three wells were exposed to full osteogenic differentiation medium (DMEM high glucose, 100nM dexamethasone, 50μg/mL 2-phospho-L-ascorbic acid trisodium salt, 10mM β-Glycerophosphate disodium salt hydrate, 10% FBS and 0.1x penicillin/streptomycin). Three wells remained in growth media as controls. Media was changed three times per week and cells were cultured for 5 weeks. Osteogenic potential was identified by Alizarin Red S staining (see Supplementary Methods).

*Chondrogenesis.* 3D cell spheroids were formed by seeding 100,000 cells/well of each cell type in a 96-well U-bottom shaped culture plate and culturing them for 48 hours in growth media. Three spheroids were kept in growth media as controls and three spheroids were exposed to full chondrogenic medium (Biological Industries MSCgo™ Chondrogenic XF Basal Medium with Chondrogenic XF Supplement Mix). Chondrogenic potential was evaluated by Alcian blue staining (see Supplementary Methods).

### Protein measurements

Cells were harvested for protein collection on days 0 and 14 of adipogenic differentiation. Proteins of interest were detected by Simple Western Assay on the WES (ProteinSimple, Santa Clara, CA). See Supplementary methods for details.

### Statistical analysis

Data management and analysis were completed in SAS v 9.4 (Cary, NC) and GraphPad Prism version 9.1.1 for macOS (GraphPad Software, San Diego, CA). Baseline differences between the four groups and growth success rates between groups and lineages were tested by chi-square test for non-numeric variables. Differences between the four groups in baseline characteristics and cell populations were tested using a 2 x 2 general linear model with Type III sum of squares to account for differing sample sizes between the groups. Tukey’s HSD post-hoc test was used to determine group differences when the global F-statistic reached significance. Skewed variables (>1.0) underwent log transformation before statistical testing, but data included in figures and tables is presented untransformed for clarity. Skewed variables included: ATM & Monocytes (% CD45^POS^), basophils (% CD45^POS^), CD31^NEG^/CD29^POS^ (% CD45^NEG^). Two-way ANOVA was used to test the effects of lineage and time on cell proliferation and one-way ANOVA to test lineage differences in ORO accumulation and protein content, with Bonferroni correction. Group differences were not tested for proliferation, ORO or protein content due to limited sample size and lack of statistical power, thus those analyses are pooled for sex and age. Sample sizes are included in the figure and table legends. The standard convention of controlling the type 1 error rate at 0.05 was used. Data are reported as means ± SEM or n (%), unless otherwise specified.

## Results

### Subject characteristics

By design, older and younger groups differed by age (**Table 1**). BMI did not differ between groups, although height and fat-free mass were greater in men and percent fat mass greater in women, irrespective of age. As expected, premenopausal women had higher estradiol levels and postmenopausal higher FSH. Testosterone was not different in older and younger men. Although within the normal range, glucose was higher in the older compared to younger groups for both sexes. Insulin did not differ between groups.

**Table 1.** Participant characteristics.

|  | Young Female<br>(n=13) | Older Female<br>(n=15) | Young Male<br>(n=12) | Older Male<br>(n=15) | p-value |
| --- | --- | --- | --- | --- | --- |
| Ethnicity, n (%) |  |  |  |  | 0.034 |
| Not Hispanic | 9 (69) | 13 (87) | 12 (100) | 15 (100) |  |
| Hispanic | 4 (31) | 2 (13) | 0 (0) | 0 (0) |  |
| Race, n (%) |  |  |  |  | 0.18 |
| White | 13 (100) | 14 (93) | 10 (83) | 14 (93) |  |
| Asian | 0 (0) | 0 (0) | 2 (17) | 0 (0) |  |
| Black/African American | 0 (0) | 0 (0) | 0 (0) | 1 (7) |  |
| More than one race | 0 (0) | 1 (7) | 0 (0) | 0 (0) |  |
| Age, yrs | 38.6 ± 6.7 | 61.5 ± 6.9* | 28.0 ± 3.2<sup>#†</sup> | 60.9 ± 6.2*<sup>†</sup> | <0.001 |
| Height, cm | 163 ± 5 | 164 ± 7 | 178 ± 6<sup>#†</sup> | 179 ± 7<sup>#†</sup> | <0.001 |
| Weight, kg | 69.0 ± 8.1 | 74.7 ± 13.8 | 77.7 ± 10.2 | 89.1 ± 15.2<sup>#†</sup> | 0.001 |
| BMI, kg/m<sup>2</sup> | 26.2 ± 3.7 | 27.6 ± 4.6 | 24.7 ± 3.2 | 27.7 ± 3.0 | 0.13 |
| Fat-free Mass, kg | 43.0 ± 4.6 | 44.5 ± 8.5 | 59.0 ± 6.1<sup>#†</sup> | 63.8 ± 8.7<sup>#†</sup> | <0.001 |
| Fat Mass, kg | 25.6 ± 5.6 | 30.3 ± 8.9 | 18.7 ± 5.9<sup>†</sup> | 25.3 ± 7.7 | 0.002 |
| Body Fat, % | 37.1 ± 5.1 | 40.2 ± 7.0 | 23.8 ± 4.6<sup>#†</sup> | 27.9 ± 4.5<sup>#†</sup> | <0.001 |
| Estradiol, pmol/L | 675 ± 554 | 55 ± 22 | ND | ND | <0.001 |
| FSH, mIU/mL | 6 ± 4 | 79 ± 31 | ND | ND | <0.001 |
| Testosterone, nmol/L | ND | ND | 18.5 ± 4.2 | 16.1 ± 4.3 | 0.17 |
| Glucose, mmol/L | 4.7 ± 0.3 | 5.5 ± 0.7* | 4.7 ± 0.4<sup>†</sup> | 5.1 ± 0.4 | 0.001 |
| Insulin, uIU/mL | 7 ± 3 | 6 ± 2 | 4 ± 2 | 5 ± 3 | 0.12 |

Data are mean ± SD unless otherwise stated. FSH, follicle stimulating hormone; BMI, body mass index; ND, measurement not done. Chi squared used for categorical variables. 2 x 2 General Linear Model with Tukey post hoc adjustment for group differences. \*p<0.05 vs same sex, # p<0.05 vs same age, † vs opposite age and sex. For conversion to conventional unit, divide by 3.671 for estradiol (pg/mL), 0.0347 for testosterone (ng/dL), and 0.0555 for glucose (mg/dL).

### Flow cytometry identification of cell populations

The results of SVF cell population analysis via flow cytometry are presented in **Figure 2**. Mesenchymal lineage (CD45^NEG^) cells made up the majority of the SVF cells in all groups except for young women, although there was a main effect of sex indicating that men had more CD45 negative cells overall (**Figure 2A**). Within the mesenchymal fraction of cells, older women had more conventional lineage preadipocytes as identified as CD45^NEG^/CD31^NEG^/CD29^POS^ than any other group (**Figure 2B**). Women had more hematopoietic cells as a fraction of all live SVF cells, with young women having significantly more than both age groups of men (**Figure 2C**).

**Figure 2.**
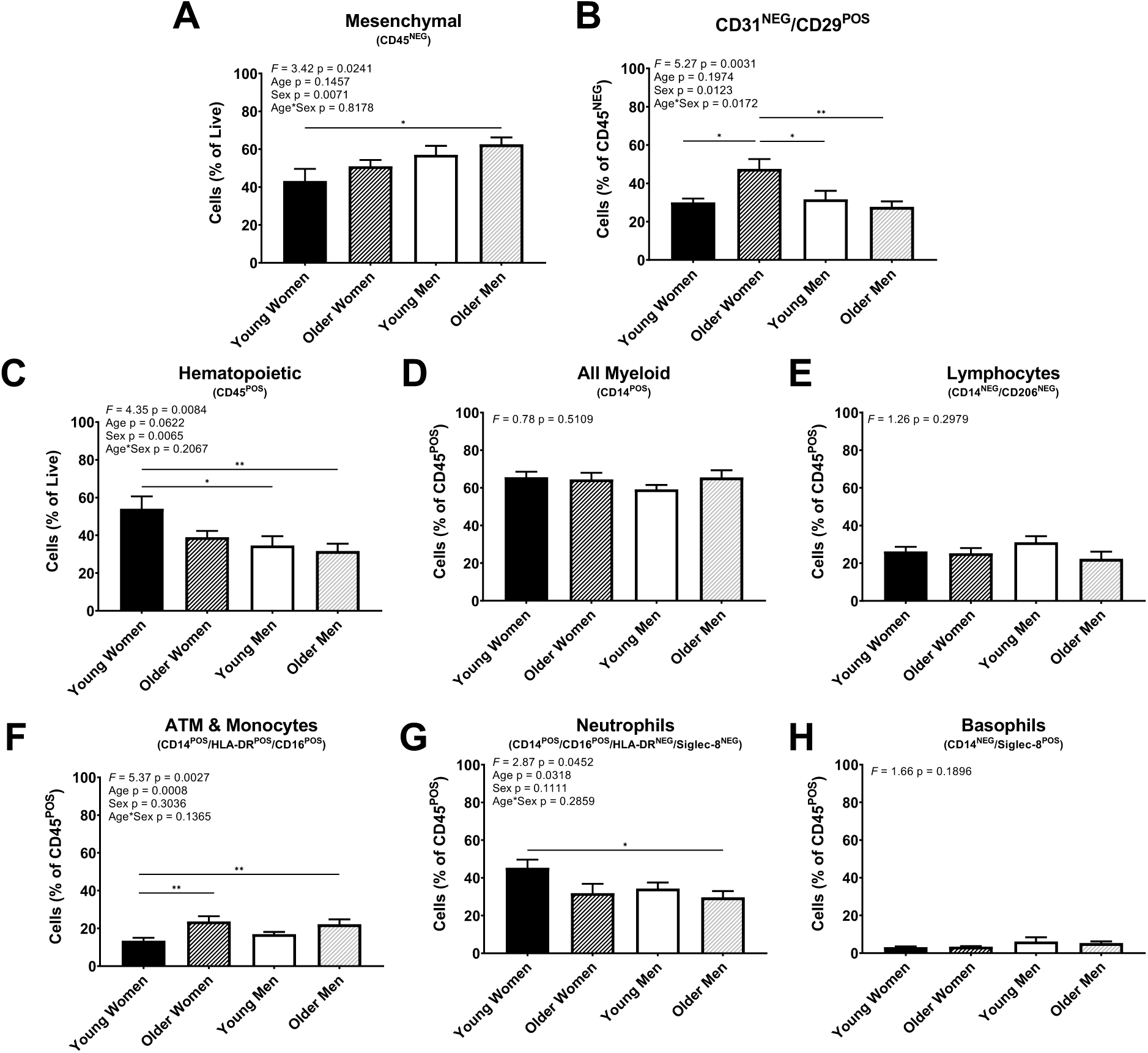
Flow cytometry identification of stromal vascular fraction cell populations. ATM, adipose tissue macrophage. Analyses: 2 x 2 general linear model with Tukey post-hoc testing. *p<0.05, **p<0.01, ***p<0.001, start and end of line indicates group differences. Sample sizes identical to Table 1.

The proportion of myeloid (CD14^POS^) and lymphoid (CD14^NEG^) lineage cells within the hematopoietic fraction was similar between groups (**Figures 2D & E**). A main effect of age indicated a greater proportion of ATM & monocytes in the SVF of older men and women, with younger women having fewer than both the older women and men (**Figure 2F**). The proportion of neutrophils was greatest in young women (**Figure 2G**) while the proportion of basophils was similar between groups (**Figure 2H**).

Flow analysis of blood samples revealed the presence of all hematopoietic lineage cells except for macrophages and did not identify any mesenchymal lineage cells (**Supplementary Table 1**).

### Culture of hematopoietic lineage cells from adipose tissue SVF

All types of hematopoietic cells isolated from adipose tissue SVF, but none from the circulation, were capable of plastic adherent growth after retrieval from fibrin (**Table 2**). Donor samples of mesenchymal lineage cells grew significantly more often than any of the hematopoietic lineage cells (Ξ^2^, 15.9, p=0.0012).

**Table 2.** Successful *in vitro* proliferation of mesenchymal and hematopoietic lineage cells isolated from adipose tissue and circulation.

| Cell Type | All | Young Female | Older Female | Young Male | Older Male | Number Sorted (x10<sup>6</sup>) |
| --- | --- | --- | --- | --- | --- | --- |
| Stromal Vascular Fraction |  |  |  |  |  |  |
| Mesenchymal Lineage | 49 (100) | 13 (100) | 13 (100) | 10 (100) | 13 (100) | 0.161 ± 0.019 |
| ATM/Monocytes | 36 (73.5) | 8 (61.5) | 11 (84.6) | 7 (70) | 10 (76.9) | 0.175 ± 0.026 |
| Neutrophils | 37 (78.7) | 9 (69.2) | 10 (83.3) | 7 (77.8) | 11 (84.6) | 0.180 ± 0.030 |
| Lymphocytes | 34 (73.9) | 11 (84.6) | 11 (84.6) | 3 (37.5) | 9 (69.2) | 0.148 ± 0.023 |
| Blood Cells (n=12) | 0 (0) | 0 (0) | 0 (0) | 0 (0) | 0 (0) | 0.325 ± 0.082 |

Data are number of samples that proliferated successfully on plastic for each group/cell type (n (%)) and number of cells sorted (mean ± SEM). ATM, adipose tissue macrophage. ATM/Monocytes, Neutrophils, Lymphocytes were grown on plastic after 5-7 days in 3D fibrin clot.

Proliferation and differentiation capacity was measured in a subset of samples. Because of the limited sample size, no age or sex comparisons were completed. Importantly, samples from both age groups and sexes were included in all experiments. Typically, if the cells underwent successful proliferation, they also successfully underwent some extent of mesenchymal differentiation. The number of viable cells (indicative of proliferation), increased over time, with no difference between lineages (**Figure 3A**). Cells of both mesenchymal and hematopoietic lineages demonstrated multilineage potential, differentiating into adipocytes, osteocytes, but not robustly into chondrocytes (**Figure 3B**). Specifically, there was no difference in adipogenic differentiation capacity between lineages as indicated by Oil red O quantification (**Figure 3C**) and content of major adipocyte proteins (**Supplementary Figure 3**). Thus, after isolation from adipose tissue, hematopoietic lineage cells cultured in a 3D fibrin clot can undergo hematopoietic-to-mesenchymal like transition confirmed by multilineage differentiation potential.

**Figure 3.**
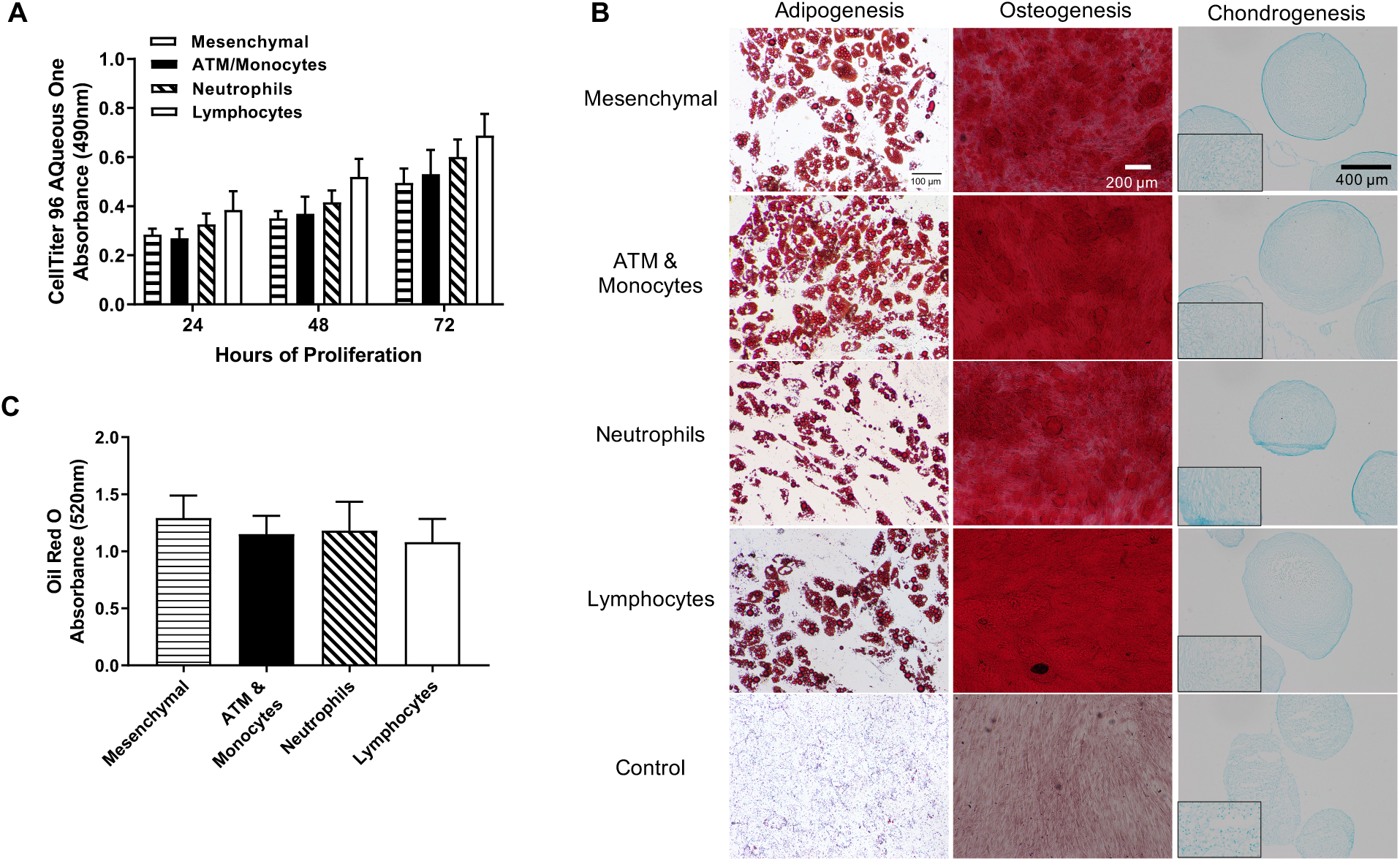
Proliferation and differentiation did not differ between adipocyte precursor lineages. **(A)** Cell proliferation was measured at 24, 48, and 72 hours after plating. Analysis: two-way ANOVA with significant main effect of time (p<0.0001) but not cell lineage (p=0.2). *** p<0.001, **** p<0.0001 **(B)** Representative images of each cell lineage after undergoing adipogenic, chondrogenic or osteogenic differentiation as indicated by positive staining for Oil Red O, Alizarin Red, or Alcian Blue, respectively. Undifferentiated cells are from same donor with identical staining to differentiated samples. Image magnifications: adipogenesis 10X, osteogenesis 4X, chondrogenesis 4X (with 10X inset) **(C)** Adipogenic differentiation capacity was not different between lineages. One way ANOVA F=0.194, p=0.89. Sample sizes: Mesenchymal n=10, ATM/Monocytes n=7, Neutrophils n=8, Lymphocytes n=8. (Use color in print version)

## Discussion

Studies utilizing murine lineage tracing strategies and single cell sequencing demonstrate that adipocyte precursors are comprised of a heterogeneous cellular population.(9, 18) Here we demonstrate that, indeed, hematopoietic lineage cells isolated from the adipose tissue stroma of human donors are capable of mesenchymal transition and multilineage differentiation. This supports our previous findings that some adipocytes in the major white fat depots of mice and humans arise from a formerly unappreciated source of adipocyte progenitors, the hematopoietic lineage.(6, 10, 11, 13)

The lack of a novel biomarker to identify BMDAs at the mature adipocyte level in humans is a barrier to advancement in the field. Murine studies provide a model in which labeled bone marrow transplantation, lineage tracing, or ablation can be utilized to identify cellular lineage and mechanistic underpinnings of cell development and consequences to their presence or absence. Clinical studies of cellular lineage are much more challenging. To date, the only studies of BMDAs in humans were conducted in a clinical population of hematopoietic stem cell transplant (HSCT) patients in which cellular lineage can be identified using donor/host DNA chimerism.(6, 14) Although a robust and reliable methodology, this model and patient population does not provide the tools necessary to understand the true clinical relevance of BMDAs and their potential role in human health and disease.

The 1960s brought evidence of immune cells in adipose tissue.(19, 20) Recent studies highlight the complexity of the adipose tissue immune cell environment.(21, 22) Moreover, adipose tissue is an extramedullary reservoir for hematopoietic stem and progenitor cells, albeit to a much lower extent than bone marrow.(23) Gerard et al. first reported hematopoietic lineage cells undergoing differentiation into mesenchymal lineage cells,(24) a finding we confirmed in hematopoietic lineage cells isolated from adipose tissue of mice.(13) Here we utilized flow cytometry to identify mesenchymal and hematopoietic lineage cells in the SVF of human adipose tissue and adapted our previously published methodology (13) to develop a human primary cell culture model of BMDAs, which will facilitate translational studies of BMDAs.

Previous murine studies identified the hematopoietic lineage source of BMDAs to be myeloid in origin.(11, 15) Corroborating those findings, we found myeloid lineage cells (i.e., macrophages, monocytes and neutrophils) were capable of adipogenic differentiation. However, in our model, cells of lymphoid origin also displayed adipogenic capacity, although this tended to happen less reliably. This could be due to differences in the culture methodology, or the sorting strategies used to identify lymphoid populations between the murine and human studies. It is also possible that there is more lineage plasticity in human compared to murine lymphoid cells. Further discrimination of lymphoid subpopulations and their ability to contribute to BMDA production is needed.

Cells from both mesenchymal and hematopoietic lineages were capable of proliferation and differentiation, although non-mesenchymal lineage cells did not successfully proliferate *in vitro* as consistently as those of the mesenchymal lineage. Although not statistically significant, hematopoietic lineage cells isolated from older men and women tended to grow more successfully (∼81% of the time) than those from younger donors (∼68% of the time). Murine studies demonstrate BMDAs are present at greater proportions with age or time since transplant,(6, 11) thus, it is possible that age plays a role in the success of primary BMDAs grown in culture. Additional studies are warranted to investigate the effect of age.

Whether hematopoietic cells commit to trafficking to adipose tissue and undergoing mesenchymal transition before they leave the bone marrow niche, or it is a local environmental factor (e.g., hormone, cytokine, physical force) that initiates this unexpected differentiation pathway is unclear. We found hematopoietic lineage cells isolated from circulating blood samples were not capable of mesenchymal-like transition, despite identical methodology to SVF-derived hematopoietic lineage cells. This aligns with our previous murine studies demonstrating hematopoietic lineage cells isolated from circulation, bone marrow, peritoneal fluid, liver or lung were not capable of mesenchymal transition using similar methodology.(13) These data support the hypothesis that an adipose tissue microenvironmental factor(s) is responsible for the hematopoietic-to-mesenchymal transition of some hematopoietic lineage cells located in the adipose tissue. More research is needed to discover the mechanism responsible for this transition.

We found older women had more conventional mesenchymal lineage preadipocytes than any other group. The presence of more adipocyte progenitors may prime the abdominal depot for adipogenesis and be an underpinning of the increased abdominal adiposity exhibited by women over the menopausal transition.(25) Unexpectedly, we found young women had the greatest proportion of hematopoietic lineage cells within the SVF. This seemed to be driven by the accumulation of neutrophils. Notably, the infiltration of neutrophils into adipose tissue is recognized as an early step in the development of obesity-associated adipose tissue inflammation, producing chemokines and cytokines that lead to macrophage infiltration.(26, 27) The BMI range of the young women in this study was 21-34 kg/m^2^, with the mean BMI in the overweight category. The elevated neutrophil infiltration could be reflective of the initiation of adipose tissue inflammation. Furthermore, older women had a greater proportion of ATM and monocytes than any other group. Although cross-sectional, this could reflect the subsequent step in the inflammatory cascade to the elevated neutrophils observed in the premenopausal women. Our work in murine models and the *in vitro* studies here suggests that ATMs are BMDA precursors.(11, 13) Thus, it is possible that the presence of more ATMs in postmenopausal women results in greater production of BMDAs. This hypothesis is consistent with our previous finding that ovariectomy, a surgical model of menopause, increased production of BMDAs.(12)

This study has limitations. First, we only investigated three broadly identified hematopoietic populations. We appreciate that the hematopoietic lineage is much more complex and future studies should examine these details. Furthermore, the purity of our sorting strategy for the hematopoietic lineage populations was <100%. Thus, it is possible some mesenchymal lineage cells contaminated the hematopoietic lineage cell culture studies. We did not observe any indication of proliferative senescence (e.g., reduced proliferative or differentiation capacity, increased cell size, flattening of cells, or irregular cell shape), which would have been probable with the number of population doublings necessary for a few mesenchymal lineage cells to usurp the hematopoietic lineage cultures.(28, 29) We did not investigate whether additional *in vivo* factors such as hormone environment or changes in tissue structure or microenvironment play a role in the development of BMDAs, but this is an important future direction.

In conclusion, some hematopoietic lineage cells located in the SVF of adipose tissue of humans are capable of adipogenic differentiation. This further confirms the clinical translation of BMDAs and the importance of understanding their role in maintaining or perturbing metabolic homeostasis. We also provide a novel primary cell culture model for BMDAs that can be utilized for translational mechanistic studies.

## Acknowledgments

Thank you to the participants who volunteered for the study. We would also like to thank Rebecca Benson, PA-C for completing the adipose tissue biopsies, Nicholas Williams for his assistance with the WES assays and the Morphology Phenotyping Histology Core Lab at the Charles C. Gates Center for Regenerative Medicine for their assistance. Funding support came from: NIH grants K01 DK109053, UL1 TR002535, P30 CA046934, DK048520, the Boettcher Foundation, and the Ludeman Family Center for Women’s Health Research.

## Supplementary Information

### Supplementary Methods

#### Body composition

DXA intra-instrument coefficients of variation (CV) completed on women and men, aged 23-86 y, for fat-free mass and fat mass were 0.5 ± 0.3%, and 2.0 ± 1.7%, respectively. Scans were completed by trained and experienced technicians and reviewed by the primary investigator to ensure correct acquisition and analysis.

#### Circulating factors

Estradiol (E2) and FSH, in samples from females only, were measured by chemiluminescent immunoassay (Beckman Coulter). The respective within- and between-day CVs and sensitivities for assays were: E2, 4.3%, 8.2%, 36.7 pmol/L; FSH, 1.8%, 3.8%, 0.11 mIU/mL; Testosterone, 2.1%, 5.1%, 0.59 nmol/L; glucose, 0.67%, 1.44%, 10 mg/dL; Insulin, 1.6%, 2.8%, 0.5 uIU/mL.

#### Oil Red O Staining

Six wells of each cell type were grown in 96 well plates seeded at 2,000 cells/well. Once confluent, three wells remained in growth media to serve as undifferentiated controls and three wells switched to differentiation media. On day 14 of differentiation (or control), media was removed and cells were rinsed gently with 150µL phosphate buffered saline (PBS) followed by fixation in 100µL 4% paraformaldehyde (PFA) at 4°C. After 30 minutes, cells were rinsed with 150µL PBS followed by 100µL 60% isopropanol. After removing the isopropanol, 75µL Oil red O (ORO) staining solution was added and allowed to stain for 15 minutes at room temperature. The ORO stain was saturated in 100% 2-Propanol and diluted to 60% with sterile nano-pure water immediately before use. After staining was complete, wells were quickly rinsed with 50µL of 60% isopropanol. 100µL of PBS was then added to each well and photographs were taken at 10X magnification to document differentiation capacity. ORO dye was extracted with 100% isopropanol for 10 minutes with rocking and repeated twice. Extracts were pooled and ORO absorbance measured at 520nm in triplicate.

#### Alizarin Red S Staining

After 35 days of differentiation, cells were fixed in 70% cold ethanol for 30 minutes and stained in Alizarin Red S (ARS) staining solution at room temperature for 30 minutes. Images were acquired (4X magnification) to assess osteogenic differentiation capacity.

#### Alcian Blue Staining

Chondrogenic differentiation was evaluated after 21 days of differentiation. Spheroids were fixed in 4% PFA at room temperature for 30 minutes, embedded in Optimal Cutting Temperature (OCT) compound and frozen using liquid nitrogen vapor. Sections (3 µm) (Microm HM550 Cryostat, Microm International GmbH, Walldorf, Germany) cut from the OCT-embedded samples were collected on glass slides, which were submerged in 100% ethanol for 10 seconds, washed in PBS for 5 minutes, stained with Alcian Blue for 30 minutes, and washed in running tap water for 2 minutes. They were then dehydrated for 3 minutes each through 95% alcohol (once) and 100% alcohol (twice) before being cleared in CitriSolv and mounted in synthetic resin (Permount Mounting Medium). Images were collected to evaluate chondrogenic differentiation capacity.

#### Protein measurements

Cells were harvested for protein collection on days 0 and 14 of adipogenic differentiation. Cells were rinsed two times with ice cold PBS after which cells were lysed for 15 minutes on ice (CelLytic Lysis buffer + 5µL protease inhibitor cocktail + 5µL phosphatase inhibitor cocktail per 1mL of buffer; 750µL buffer/10cm plate). Cells were scraped, sonicated 4 x 5 sec bursts (35% amplification) on ice, and centrifuged 15,000g for 10 minutes @ 4°C. The supernatant (avoiding contamination from any evident lipid layer) was transferred to a new chilled tube, with a separate aliquot saved for protein quantification by bicinchoninic acid (BCA) assay. Aliquots were stored at -80°C until use.

WES assays (separation module/detection module) were run according to the manufacturer’s instructions using 0.2mg/mL of protein unless otherwise stated. Results were normalized to total protein for each sample (total protein detection kit). Antibodies included: Acetyl-CoA Carboxylase, 1:200; Adiponectin, 1:4000; C/EBPα, 1:12.5; CD36, 1:100; Fatty Acid Binding Protein 4, 1:1500; Fatty Acid Synthase, 1:125; Leptin, 1:5, 0.4mg/mL; Lipoprotein Lipase, 1:100; Perilipin 1, 1:2000, 0.1mg/mL; PPARγ, 1:10; SREBP1, 1:25, 0.4mg/mL.

**Supplementary Table 1.** Blood cell population numbers as measured by flow cytometry.

| Cell Type |  |
| --- | --- |
| **<u>Hematopoietic Lineage</u>** |  |
| **Basophils** |  |
| Live Cells, % | 0.8 (0.6, 1.7) |
| CD45<sup>POS</sup> Cells, % | 1.6 (0.9, 2.5) |
| **Lymphocytes** |  |
| Live Cells, % | 13.9 (6.9, 33.5) |
| CD45<sup>POS</sup> Cells, % | 37.4 (9.7, 46.2) |
| **Monocytes** |  |
| Live Cells, % | 1.8 (0.3, 3.0) |
| CD45<sup>POS</sup> Cells, % | 2.6 (1.1, 4.0) |
| **Myeloid Cells** |  |
| Live Cells, % | 37.2 (14.3, 50.0) |
| CD45<sup>POS</sup> Cells, % | 47.6 (43.0, 77.2) |
| **Neutrophils** |  |
| Live Cells, % | 29.5 (10.6, 43.8) |
| CD45<sup>POS</sup> Cells, % | 41.6 (30.9, 54.3) |

Data presented as Median (25, 75<sup>th</sup> percentile).

**Supplementary Table 2.**
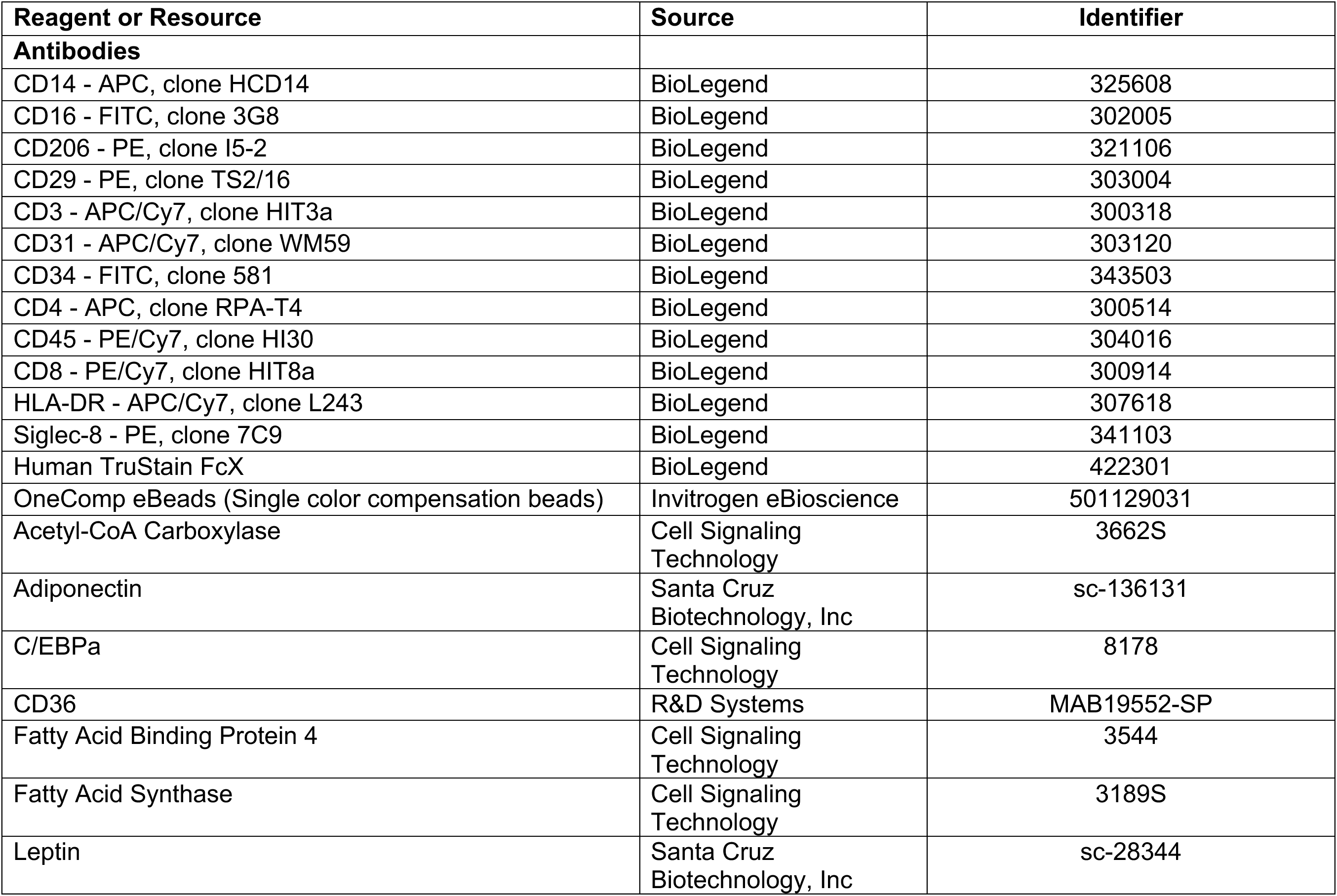
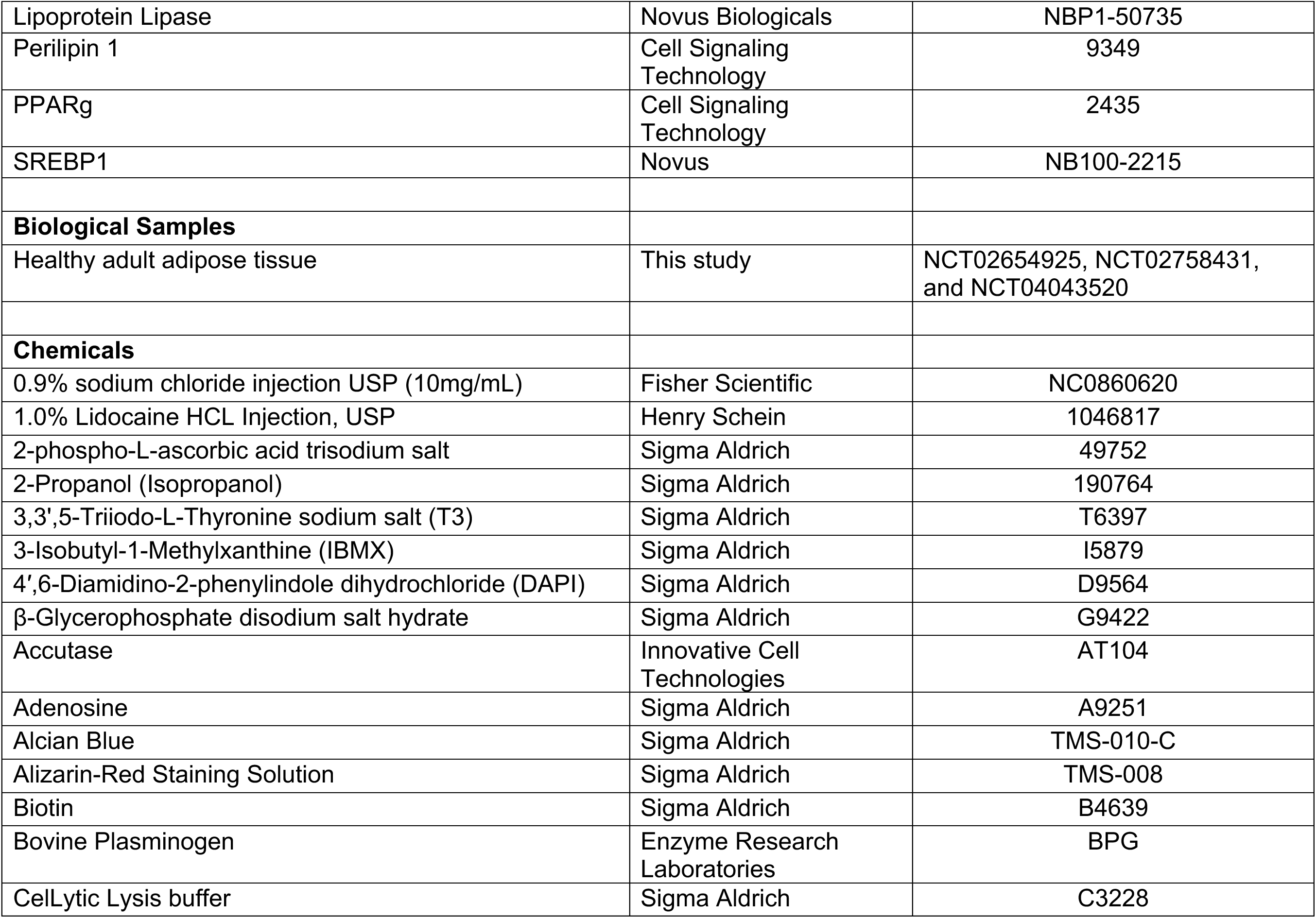
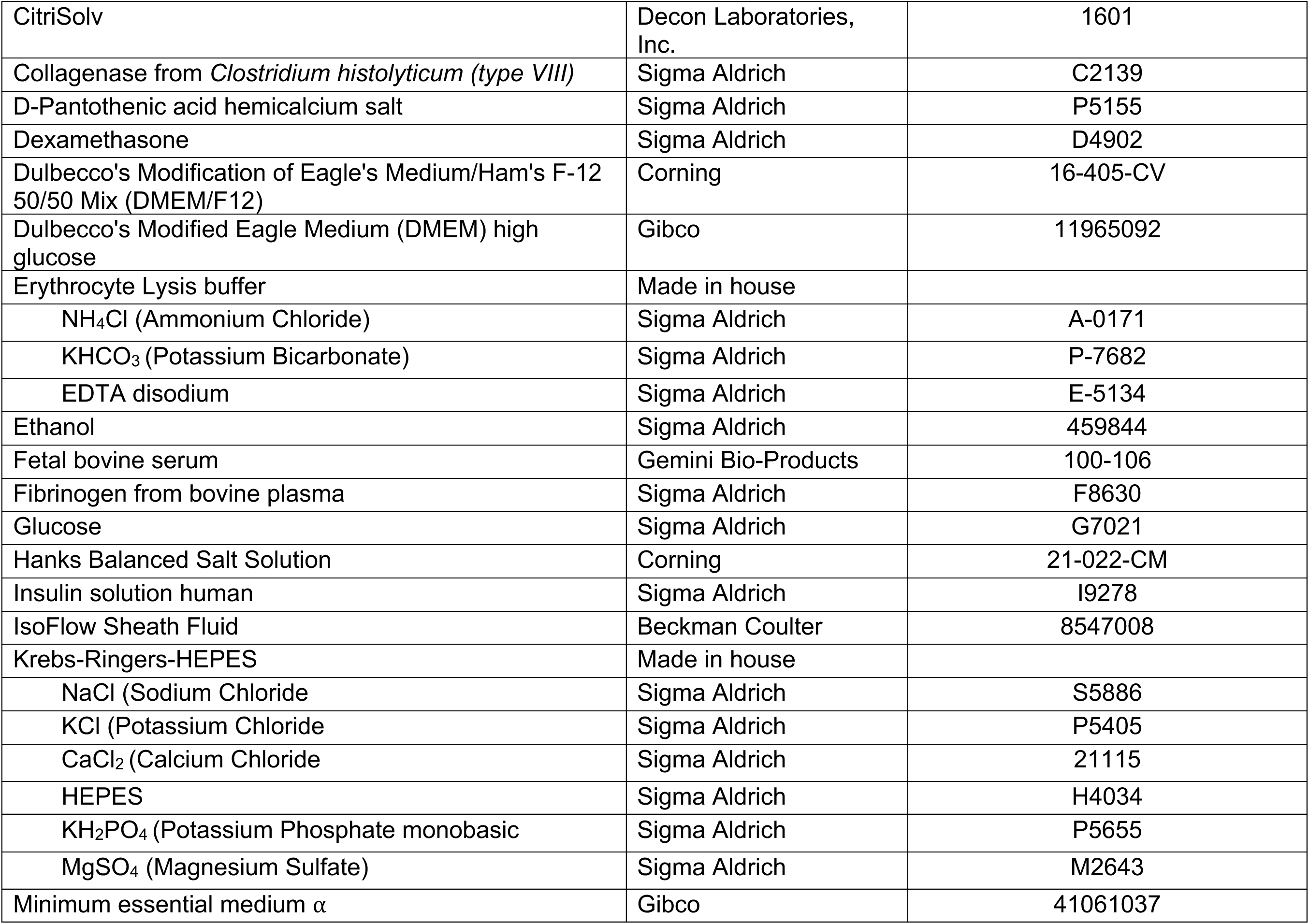
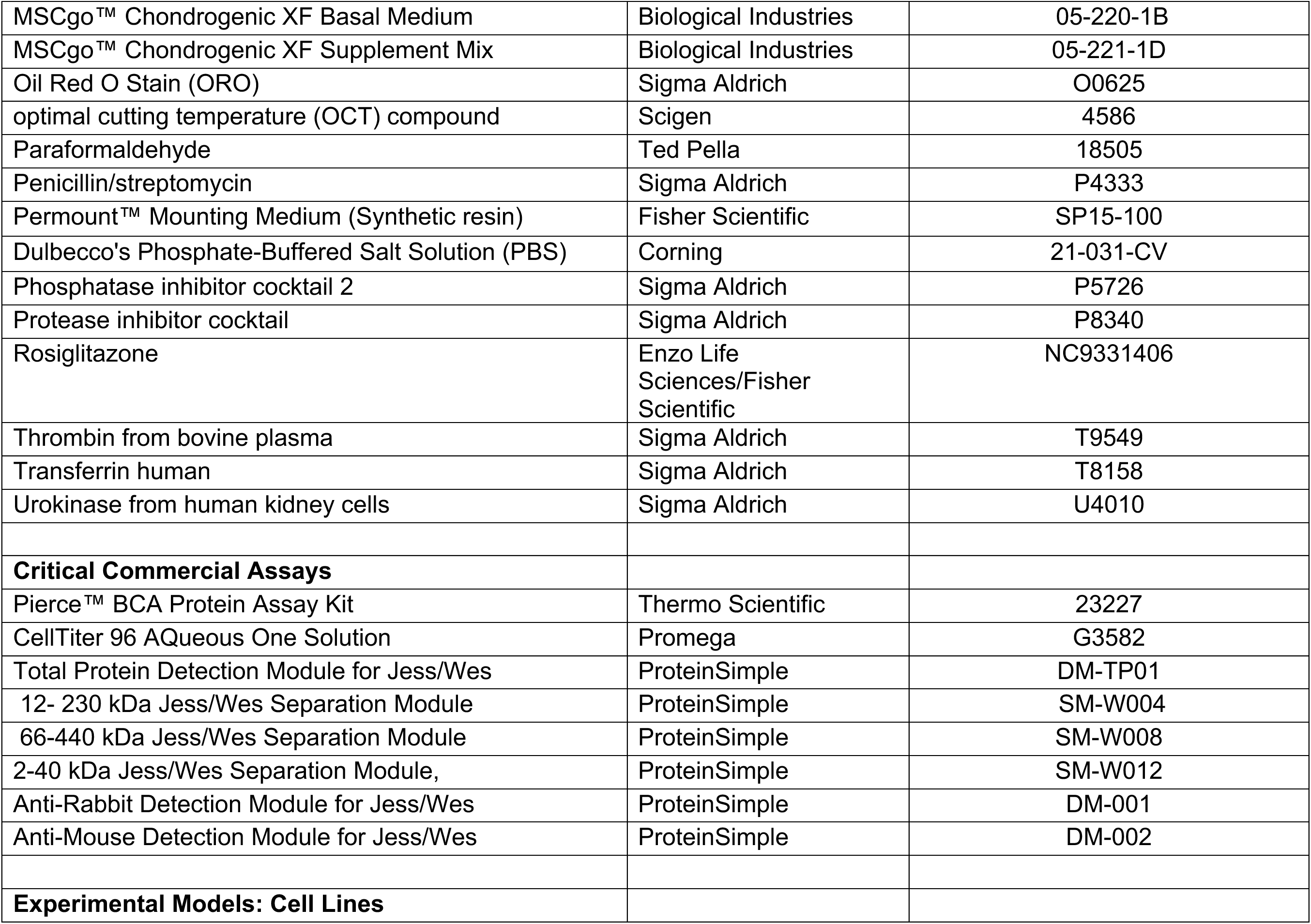
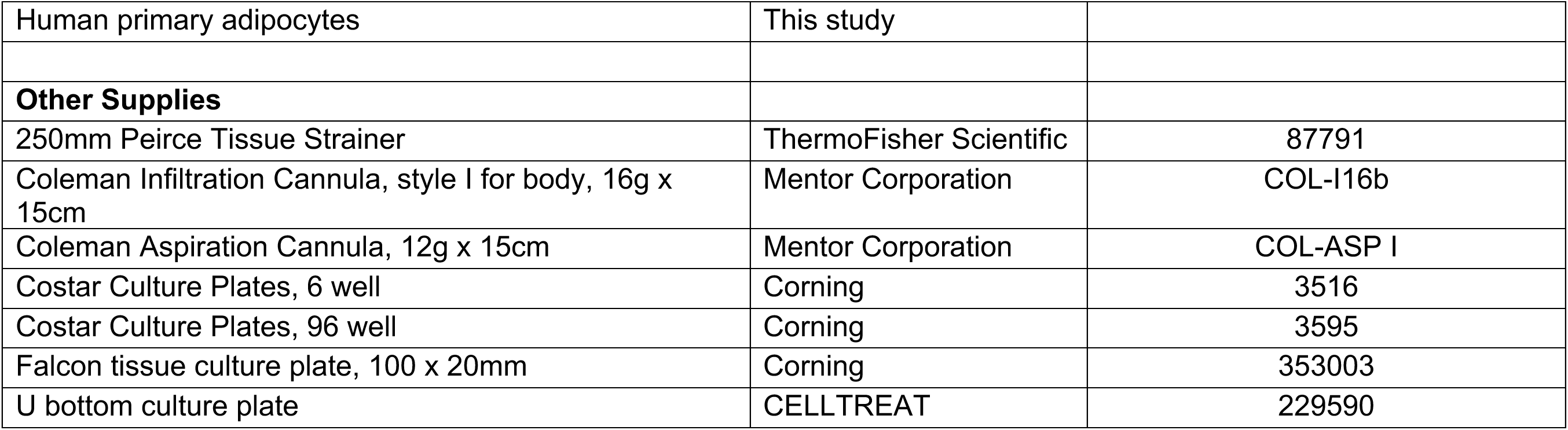
Key Resources.

**Supplementary Figure 1.**
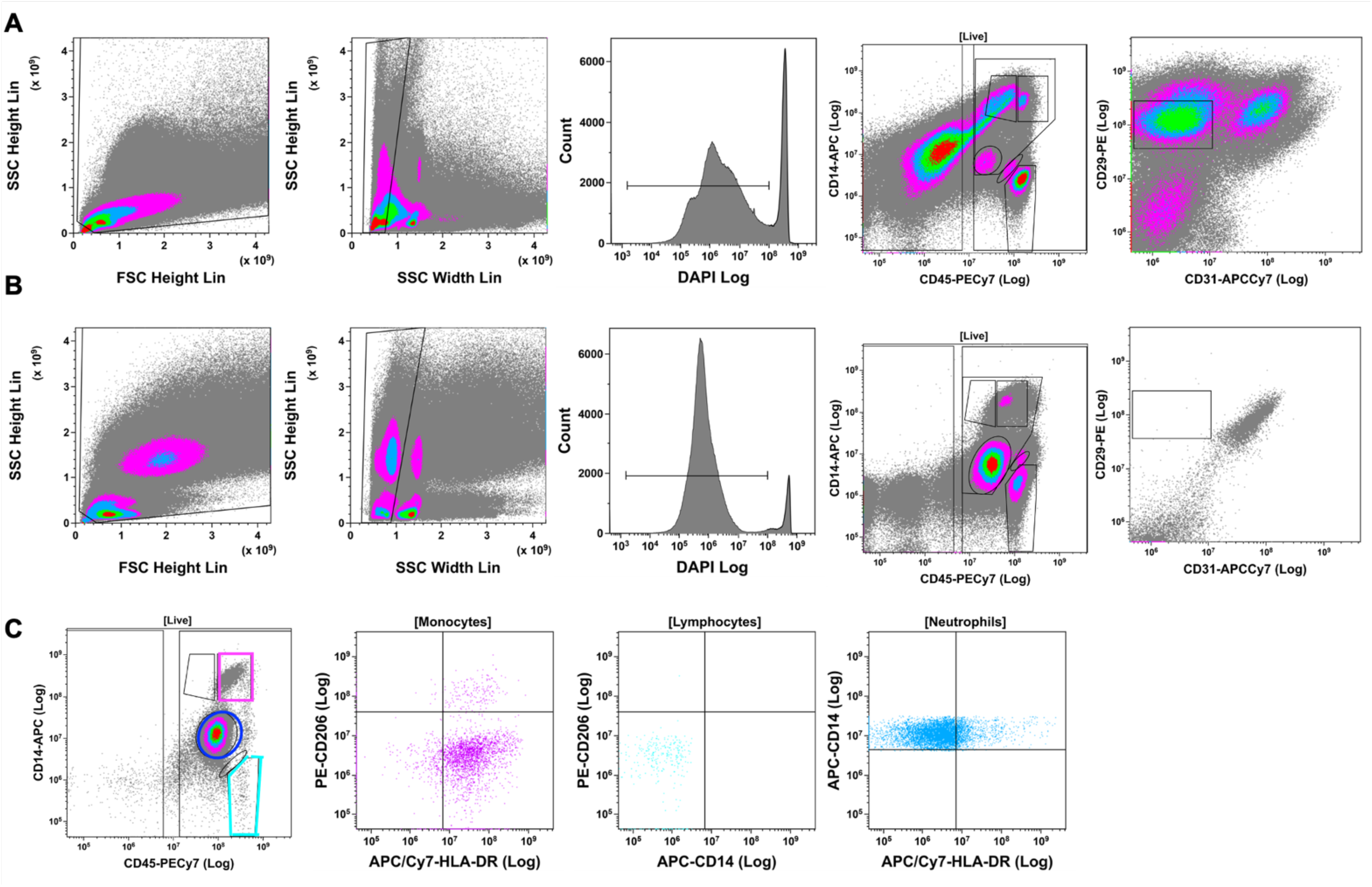
Fluorescent active cell sorting strategies for **(A)** adipose stromal vascular fraction (SVF) and **(B)** blood from the same older female donor. **(C)** Further identification of hematopoietic lineage cell populations from a representative blood sample from a second older female donor.

**Supplementary Figure 2.**
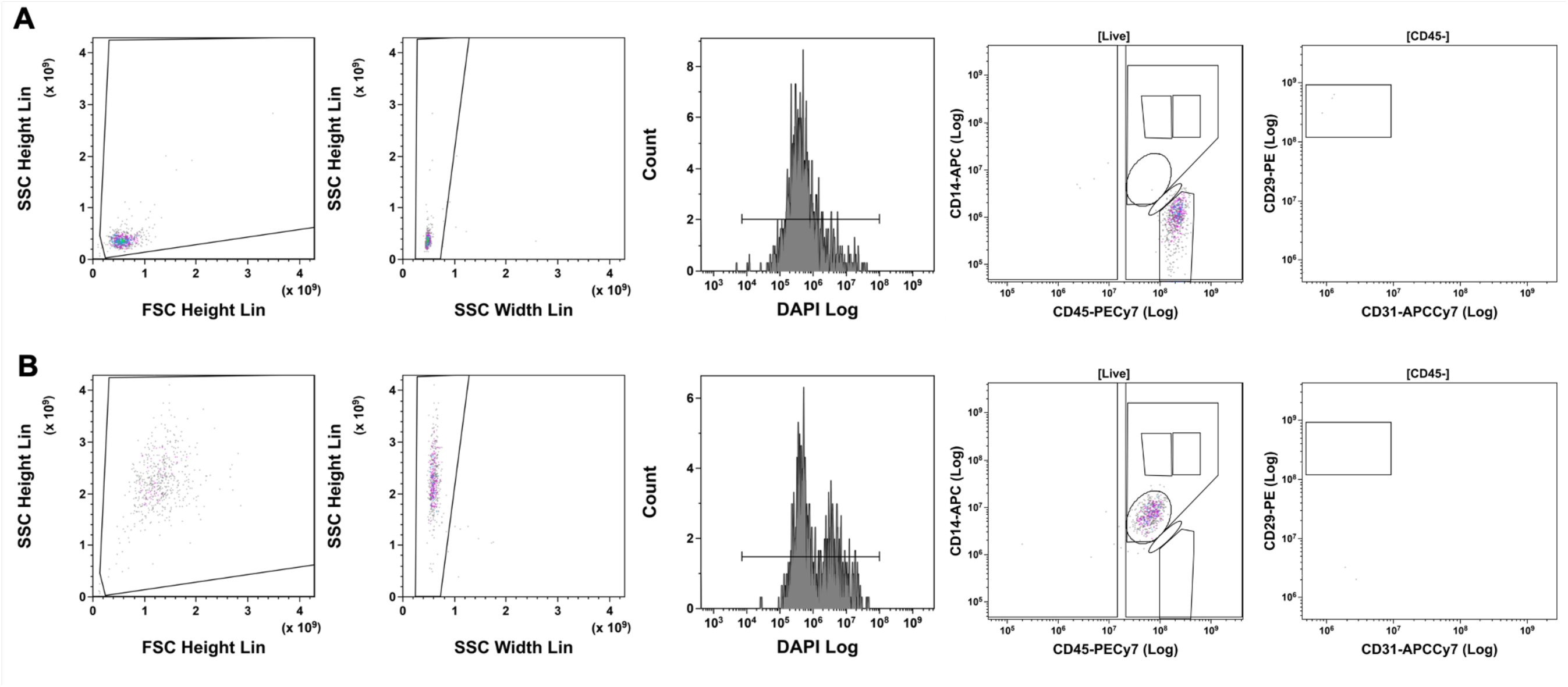
Purity of sorted cell populations. **(A)** SVF Lymphocytes. **(B)** SVF Neutrophils.

**Supplementary Figure 3.**
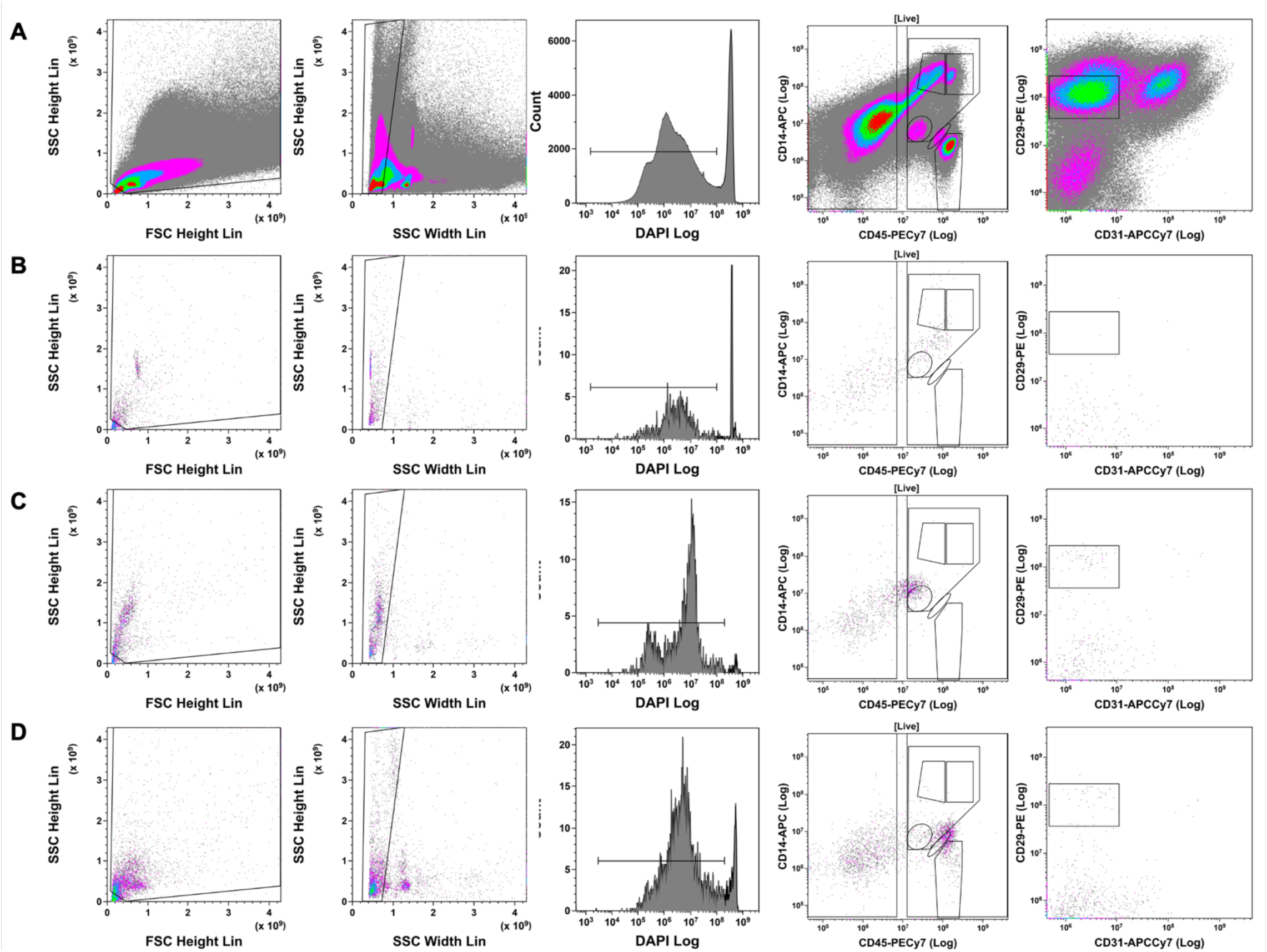
Assessment of surface markers on cells isolated from adipose SVF before and after 5-7 days of culture in 3D fibrin clot. Representative images from n=3. **(A)** Adipose SVF cell population sort before culture in fibrin clot. Adipose tissue **(B)** macrophages/monocytes **(C)** neutrophils and **(D)** lymphocytes after recovery from fibrin clot.

**Supplementary Figure 4.**
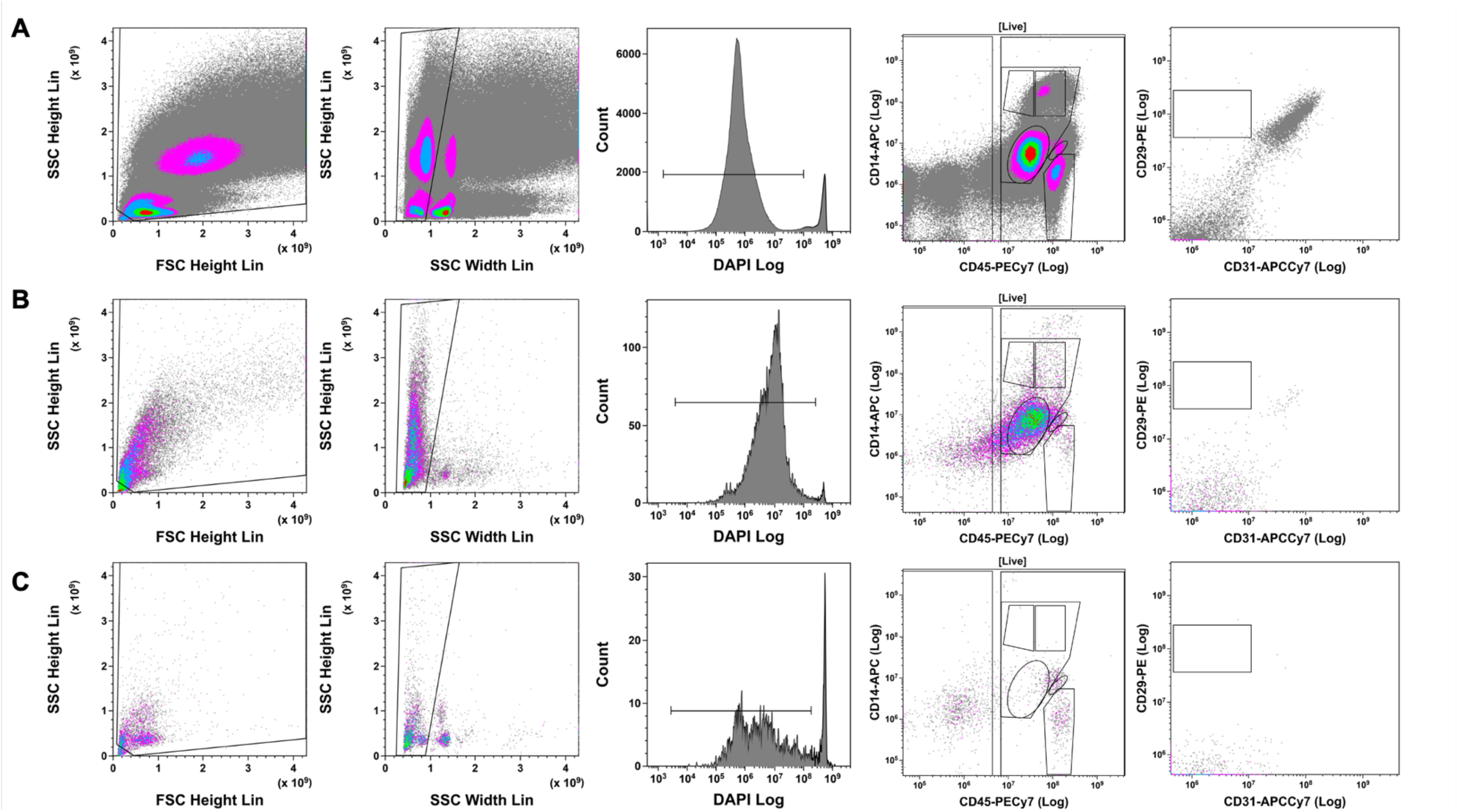
Assessment of surface markers on cells isolated from blood before and after 5-7 days of culture in 3D fibrin clot. Representative images from n=3. (**A)** Blood cell population sort before culture in fibrin clot. Blood **(B)** monocytes and neutrophils and **(c)** lymphocytes after recovery from fibrin clot.

**Supplementary Figure 5.**
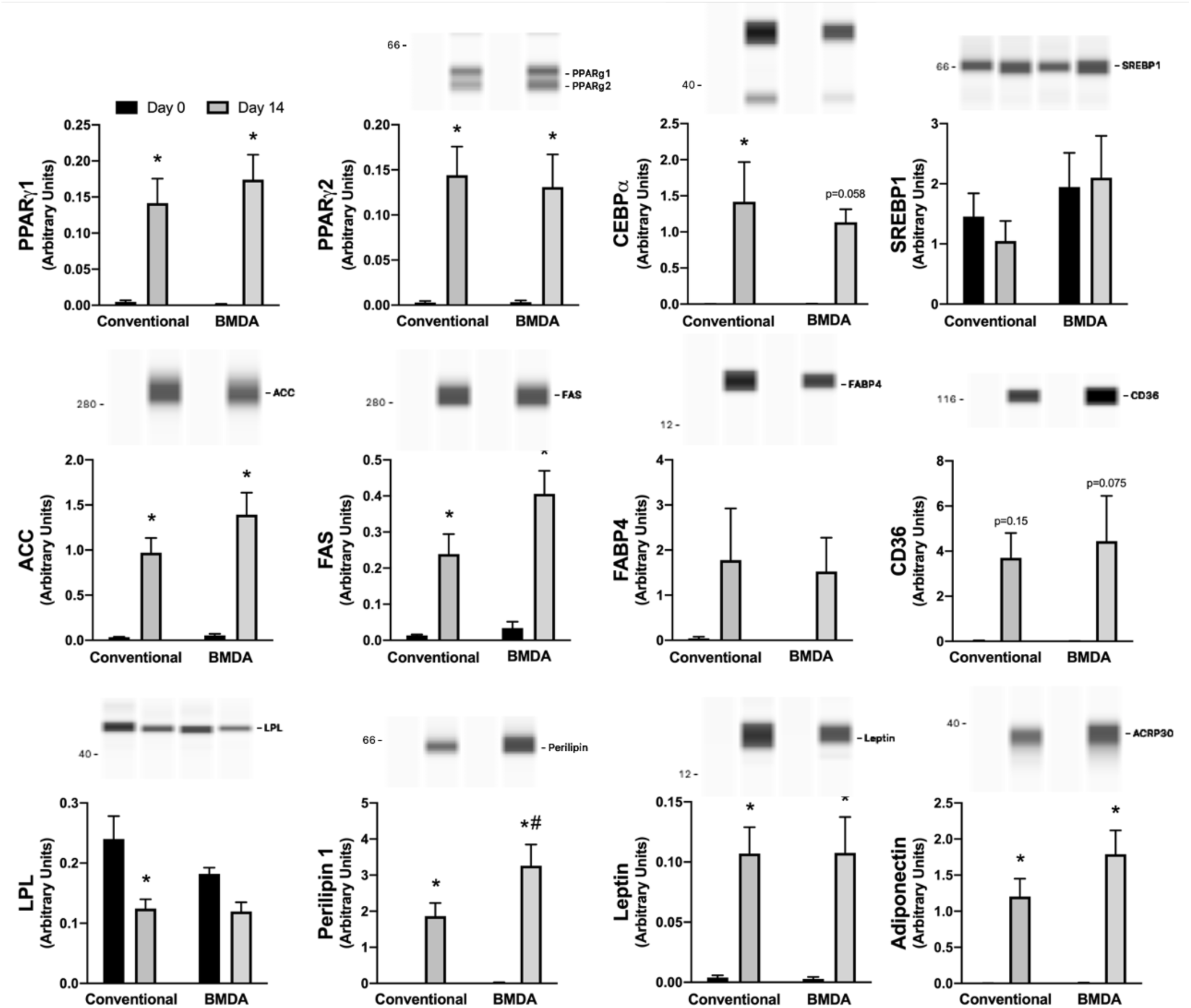
Cells from all cell linages isolated from the adipose tissue stromal vascular fraction differentiated into adipocytes as demonstrated by the presence of major adipocyte proteins. Results are normalized to total protein. One-way ANOVA, Bonferroni post hoc testing. * p<0.05 vs day 0 of same cell lineage, # p<0.05 vs same timepoint of other lineage. Samples sizes: Conventional day 0 n=5-7, BMDA day 0 n=5-8, Conventional d14 n=8-10, BMDA day 14 n=7-8. Data are mean ± SEM. Images are representative capillary images from WES. BMDA, bone marrow-derived adipocytes, a combination of lysates from adipocytes differentiated from ATM/monocyte, neutrophil and lymphocytes.

